# Production and use of triploid zebrafish for surrogate reproduction

**DOI:** 10.1101/608687

**Authors:** Roman Franěk, Tomáš Tichopád, Michaela Fučíková, Christoph Steinbach, Martin Pšenička

## Abstract

We report for first time comparison of two approaches for zebrafish triploid production using cold shock and heat shock treatment. Subsequently, produced triploid zebrafish were used as a recipients for intraperitoneal transplantation of ovarian and testicular cells originating from vas:EGFP strain in order to verify their suitability for surrogate reproduction. Heat shock treatment was far more effective evaluated as success rate of triploid production and viability in comparison to cold shock. Triploids were produced with up to 100% efficiency in particular females. As expected, all triploids were males. Subsequently, germ cells transplantation revealed that triploids are suitable surrogate hosts. Production of donor-derived sperm was achieved in 23% and 16% of triploids transplanted by testicular and ovarian cells, respectively. Success of the transplantation was confirmed by positive GFP signal detected in gonads of dissected fish and stripped sperm. Germline transmission was confirmed by fertilization tests followed by PCR analysis of embryos. Reproductive success of germline chimera triploids evaluated as fertilization rate and progeny development was comparable to control groups.

## 1 Introduction

Surrogate reproduction in fish via intraperitoneal germ cell transplantation is a promising technology for aquaculture as well preservation of endangered species when recipient species with more favorable characteristics can be used [1]. So far germ cell technologies are being applied in wide range of fish species such as, salmonids [2,3], cyprinids [4,5], Nile tilapia [6–8], medaka [9], sturgeons [10–12] or several marine fish species [13,14]. One from the main prerequisites for successful surrogate propagation is the host’s sterility ensuring no contamination of donor-derived gametes during reproduction [2] and very likely lower competition for space in gonads between endogenous germ cells and introduced exogenous germ cells resulting in higher production of donor-derived gametes.

Importance of zebrafish for surrogate reproduction is in its biological and reproductive characteristics as fast maturation or availability of transgenic lines could enable to study unknown factors affecting the success of germ cell transplantation. Moreover, zebrafish can serve as a valuable model for sterility research since several distinct methods are available for sterility induction nowadays Early ablation of primordial germ cells (PGCs) can be achieved using gene knock out approaches such as Zinc Finger Nucleases against *dead end (dnd)* gene [15], or gene knock down with *dnd* antisense morpholino oligonucleotide [16,17]. Both methods require microinjection into embryos, thus an alternative approach using immersion in vivo morpholino against *dnd* can be more convenient in case of large scale application (Wong and Zohar, 2015). Sterility was achieved via PGCs depletion in transgenic fish with artificially induced nitroreductase expression in PGCs exclusively using immersion into metronidazole enzyme which was converted into toxic metabolites and only PGCs targeted toxicity was achieved [19]. Similarly, PGCs migration was disrupted in transgenic fish which had SDF1 expression controlled by a heat shock protein. Regular event of PGCs migration is beside other mechanism guided by SDF1 gradient towards the genital ridge, however, heat treatment caused throughout expression of SDF1 resulting in migration failure and production of sterile fish [20]. A cytostatic drug such as busulfan was used successfully in combination with thermal treatment, however, this method was developed only for adult fishes, because intraperitoneal administration is necessary [21]. All abovementioned methods for sterility achievement can be regarded as relatively demanding from point of time, knowledge and equipment. Thus a simple method for sterile fish generation such as production of infertile Danio hybrids by crossing zebrafish and pearl danio is convenient from point of time and possibility to produce host by simple fertilization without additional treatment [22]. Similarly, triploid fish can be produced as hosts in surrogate reproduction technology [2,9,14]. Moreover, triploids or sterile hybrids can still be more favorable way for surrogate host production in fish species without mapped genomes or at least genes of interest not enabling use of targeted transgenic or gene silencing approaches [23], or when large scale production of sterile recipients is needed or microinjection delivery of compounds inducting sterility into the eggs is difficult due to sturdy chorion as it is known for marine fish species [24,25].

Three sets of chromosomes in artificially induced triploids cannot proceed through meiosis and gamete maturation regularly, resulting in gametogenesis arrest or aneuploid gametes production further incompatible with the proper embryonic development [26]. Triploids can be induced by pressure or temperature treatment or electric shock resulting in inhibition of second polar body extrusion. However, all abovementioned physical treatments, require equipment such as pressure chamber and thermostat respectively [27–32]. Therefore, an alternative technique using cold shock could be convenient from point of the material equipment, might has less deleterious effect on the survival but with the same efficiency of triploid induction rate as the heat shock treatment.

Method for triploid zebrafish production using heat shock treatment was published already [33]. However, we did not succeed satisfactorily using the abovementioned heat shock protocol in our laboratory. Therefore we revised the procedure for heat shock treatment and compared it with cold shock to identify an optimal condition for triploid zebrafish production with respect to achieve the highest survival and produce triploid fish. Suitability of triploid fish as surrogate recipients was tested by intraperitoneal transplantation by testicular and ovarian cells from vas:EGFP strain and subsequent production of donor-derived gametes with fertility tests confirmed by fluorescent microscopy and DNA analysis.

## 2 Material and methods

The study was conducted at the Faculty of Fisheries and Protection of Waters (FFPW), University of South Bohemia in České Budějovice, Vodňany, Czech Republic. The facility has the competence to perform experiments on animals (Act no. 246/1992 Coll., ref. number 16OZ19179/2016-17214). The methodological protocol of the current study was approved by the expert committee of the Institutional Animal Care and Use Committee of the FFPW according to the law on the protection of animals against cruelty (reference number: MSMT-6406/119/2). The study did not involve endangered or protected species. Martin Pšenička owns the certificate (CZ 00673) giving capacity to conduct and manage experiments involving animals according to section 15d paragraph 3 of Act no. 246/1992 Coll.

### 2.1 Fish and gamete collection

Mature zebrafish spawners from AB line were purchased from European Zebrafish Resource Center (Germany), vas:EGFP line was purchased from University of Liège, Belgium, were maintained in a zebrafish housing system (ZebTEC Active Blue) at 28 °C, 14L:10D photoperiod, feeding two times with Tetramin flakes and once with *Artemia* nauplii. Fish were set into the spawning chambers afternoon before the spawning (one male and one female) and separated with a barrier. On the light onset of the next day, the barrier was removed and fish were observed for oviposition. Selected fish were immediately transferred into the laboratory. Gametes for *in vitro* fertilization were obtained after anesthesia in 0.05% tricaine solution (Ethyl 3-aminobenzoate methanesulfonate). Sperm from at least 5 males was pooled together in 50 μl of Kurokura 180 solution [34], eggs were collected from each female separately and fertilized promptly. Fertilized eggs were divided into control and treated groups and were cultured at 28.5 °C constantly.

### 2.2 Triploid induction and rearing

Cold shock (CS) treatment for the given time was conducted with fertilized eggs in a plastic strainer placed in a Styrofoam box with 2 L of ice chilled water. Heat shock (HS) treatment was conducted in a plastic strainer placed in a 5L recirculated water bath with thermostat under varying conditions. Parameters used in all CS and HS trials are summarized in Table 1.

**Table 1.** Variables tested during optimization of triploid zebrafish production. mpf – minute post fertilization

| Treatment | Temperature | Duration time | Initiation time |
| --- | --- | --- | --- |
| Cold shock |  |  |  |
| CS temperature | 3, 6, 9 °C | 5 min | 1 mpf |
| CS duration | 6 °C | 5, 10, 15 min | 1 mpf |
| CS initiation time | 6 °C | 5 min | 0.5, 1 mpf |
| Heat shock |  |  |  |
| HS temperature | 41, 41.4, 42 °C | 2 min | 2 mpf |
| HS duration | 41.4 °C | 1, 2, 3, 4 min | 2 mpf |
| HS initiation | 41.4 °C | 2 min | 1, 2 mpf |

The remainder of the intact fertilized eggs from each female was kept as a non-treated control. Females (n = 7) producing eggs with fertilization rate in control below 65% were regarded as having bad quality and were excluded from results and replaced by new females. Swim-up larvae were fed by paramecium for one week, later on with *Artemia* nauplii *ad libitum* and held until the first month in an incubator in plastic boxes. Fish were then transferred into a zebrafish housing system and were kept until reaching maturity. Five females with separately fertilized eggs were used as replicates. Eggs from each female were divided into approximately same portions into 3(4) groups according to tested variables in performed treatment and one untreated control constantly held at 28.5 °C. Survival was recorder as a percentage of living embryos from number of fertilized eggs

### 2.3 Flow cytometry

Only larvae after swim-up stage were used for triploidy confirmation in trials in order to obtain results representing only viable triploids. Whole larvae (euthanized by tricaine overdosing) or later on fin clips were processed using a kit for nuclei staining CyStain UV Precise T (Sysmex Partec GmbH, Germany) according to the manufacturer’s protocol. The relative DNA content was determined using a CyFlow Ploidy Analyzer (Sysmex Partec GmbH, Germany) against samples from diploid control groups. Ten larvae were analyzed from each female in each treatment and group.

### 2.4 Transplantation

Male and female germ cell donors from vas:EGFP line were euthanized by tricaine overdosing, decapitated, the body was washed with 70% ethanol. Testis from two donors were excised aseptically. Each testis was cut into 4-6 fragments and washed several times in phosphate buffered saline (PBS) in order to remove leaking sperm. Medium for testicular tissue digestion contained 0.1% trypsin, 0.05% DNAse dissolved in PBS. Fragments were collected by a pipette and transferred into 2 mL tube with 1 mL of digestion medium and were further minced with scissors and placed on a laboratory shaker for 50 min. Digestion was terminated by addition of 1 mL L-15 and 10% FBS (v:v). The suspension was filtrated through a 30 μm nylon filter (CellTrics^®^ Sysmex, Germany) and centrifuged at 0.3 g for 10 min. The supernatant was removed and the pellet was resuspended in 40 μl L-15 with 10% FBS. Female germ cells were collected from juvenile donors (2 months, n = 5 per one transplantation trial) and digested as described for testicular cells. After centrifugation, ovarian cells suspension was washed and filtrated two times to remove excess of debris.

Triploid recipients produced by optimized HS procedure were anaesthetized at 7 dpt in 0.05% tricaine and placed on petri dish coated with 1% agar. Testicular and ovarian cell suspension was loaded into the glass capillary attached to MN-153 micromanipulator (Narishige) and FemtoJet^®^ 4x injector (Eppendorf). Triploid recipients were injected by approximately 3000-5000 of testicular cells (TC group) or 500 ovarian cells (OC group) per individual. Each transplantation trial for TC and OC groups consisted of 30 transplanted fish when triploid recipients were originating from same batch in both groups. The remainder of non-injected triploids and non HS treated diploids was kept as a control and no operation was conducted on them. Transplanted fish were left to recover in dechlorinated tap water. Survival and colonization rate of transplanted cells was monitored until adulthood. Fish were observed and photographed under a fluorescent stereomicroscope (Leica M205 FA) with fluorescent filters DAPI/FITC/TRITC (order no 10450614) or GFP (order no 10450469) equipped with camera (Leica DMC 6200).

### 2.5 Production of donor-derived gametes

All adult surviving fish were screened for positive GFP signal in their testis. GFP positive germline chimeras were set into spawning chamber afternoon (two or three males and separated one female). Males were following morning anaesthetized and sperm was collected and observed under an inverted fluorescent microscope to detect positive GFP signal in sperm. All fish producing GFP positive sperm from each transplantation trial were pooled together into one TC and one OC group and left to recover for 4 weeks.

Randomly selected fish from pooled TC and OC group (10 fish per group) were propagated by semi-natural mating and *in vitro* fertilization. Semi-natural mating was conducted in spawning chambers when one germline chimera triploid male and two AB females in reproductive condition were set together in the afternoon and separated with a barrier. Next day at the onset of light, a barrier was removed and fish were allowed to spawn for three hours. Spawned eggs were collected and the survival rate was monitored. Swim up larvae from each group were pooled and 10 individuals were selected randomly and used for PCR analysis to verify the efficiency of germline transmission. Used germline chimeras in semi-natural mating were separated and were not used for following *in vitro* fertilization. Procedure for *in vitro* fertilization was the same as described for AB line (2.1). Sperm collected from each chimeric male was stored separately in immobilizing solution. Eggs were stripped from AB females (n = 4), gently mixed, divided into approximately same portions and fertilized with sperm from chimeric triploid males individually. Control group for semi-natural and *in vitro* fertilization consisted AB females and vas:EGFP males. Survival of produced embryos was monitored. Offspring from each group were pooled together and 10 randomly selected larvae were used for PCR analysis. DNA was extracted from larvae by PureLink™ Genomic DNA Mini Kit (Invitrogen™). GFP forward primer ACGTAAACGGCCACAAGTTC, reverse primer AAGTCGTGCTGCTTCATGTG. Primers were tested for specifity. The reaction mixture for PCR contained 1 μl template cDNA, 0.5 μl forward and 0.5 μl reverse primer, 5 μl PPP Master Mix (Top-Bio) and 3 μl PCR H2O (Top-Bio). Reaction conditions were 30 cycles of 94 °C for 30 s, 58 °C for 30 s and 72 °C for 30 s. Products were analyzed on gel electrophoresis on 2% agarose gel on a UV illuminator.

### 2.6 Histology analysis

Euthanized zebrafish triploids and diploid controls were fixed overnight in the Bouin’s fixative. Samples were immersed in 70% ethanol, dehydrated and cleared in an ethanol–xylene series, embedded into paraffin blocks and cut into 4μm thick sections using a rotary microtome (Leica RM2235). Paraffin slides were stained with hematoxylin and eosin by using a staining machine (Tissue-Tek DRS 2000) according to standard procedures. Histological sections were photographed and evaluated using a microscope with mounted camera (Nikon Eclipse Ci).

### 2.7 Statistical analysis

Survival of embryos was analyzed by logistic regression with mixed effects where the treatment was set as fixed effect while females were set as random effect with different intercepts (as mentioned above, eggs in each groups were obtained from five females). Post hoc Tukey’s test was performed to find out significant differences among groups of different treatment. The effect of treatment on a number of triploids was analyzed by Friedman test where individual females were set as blocks. Differences among groups were analyzed by Post-hoc Conover test with Benjamini-Hochberg correction [35]. All analysis were performed in R software (3.5.2).

## 3 Results

### 3.1 Production of triploid recipients for surrogate reproduction

The testing of CS revealed that exposition in 6 °C water bath resulted in significantly higher triploid production and survival in comparison to CS at 3° C. Few triploids were also produced at 3 °C CS, but lower temperature was more detrimental to early embryonic development when even swim-up embryos exhibited malformations (Supplementary figure 1). Cold shock conducted at 9 °C had the lower effectiveness for triploid induction. Testing of prolonged CS duration yielded comparable fraction of detected triploids in all tested durations, however, survival rate was more favorable in 5 and 10 min lasting CS treatment. Optimized CS temperature (6 °C) and duration (5 min) were used further to test different initiation times after fertilization. Yield of triploid fish was improved significantly when CS was initiated at 30 seconds post fertilization (spf) (Figure 1A-C).

**Figure 1.**
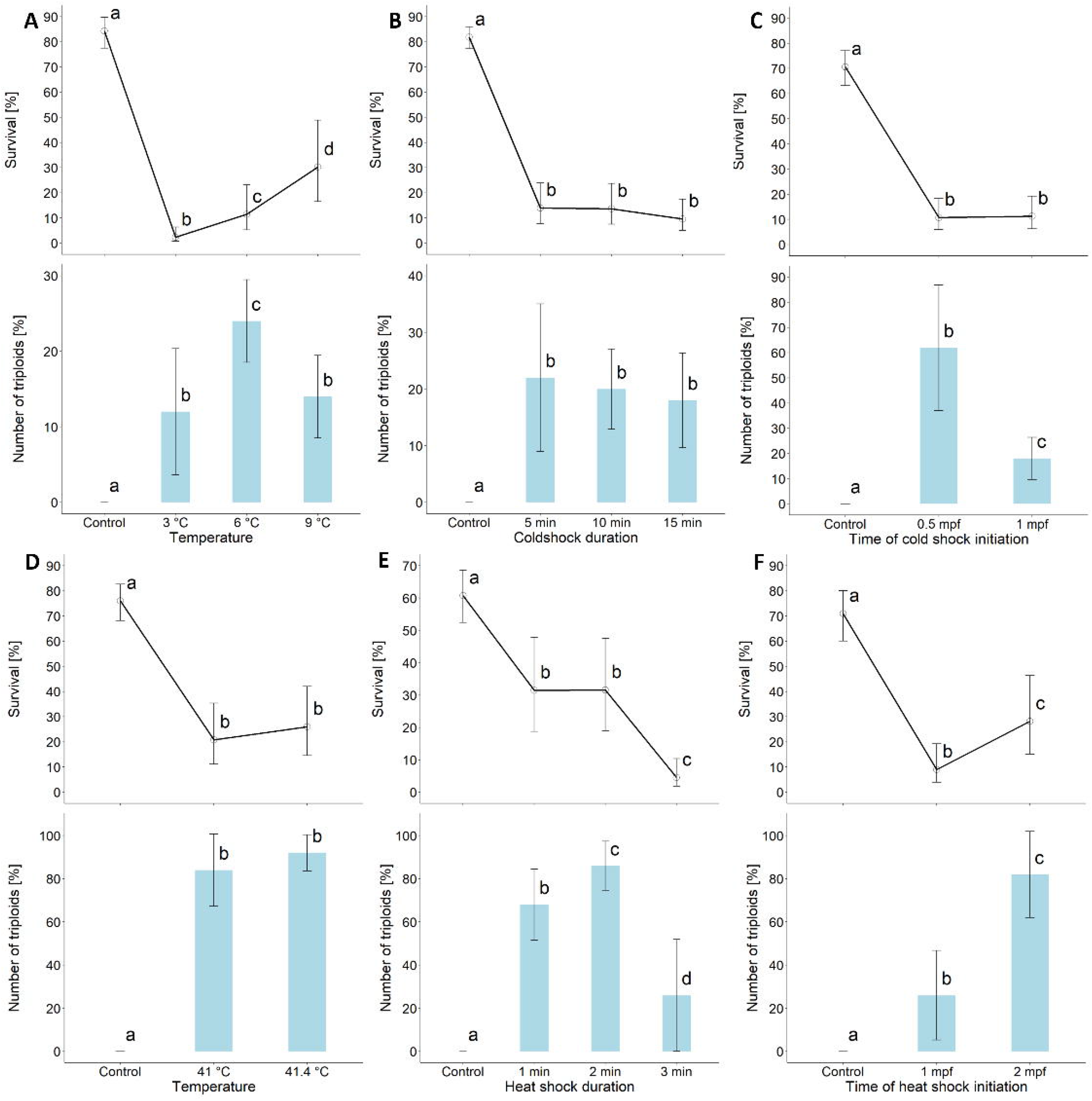
Survival and success rate of triploid induction using cold shock (A-C) and heat shock treatment (D-F). D) result from treatment at 42 °C were excluded because of total mortality of treated embryos. E) Results from heat shock duration for 4 min were excluded because of total mortality of treated embryos. Different letters above the confidence intervals (survival) or SD lines (number of triploids) indicate statistical significance (Tukey’s HSD, p < 0.05).

Triploid induction by HS treatment tested different temperatures and shock duration during the first trial. HS treatment at 42 °C was lethal for all embryos (data not shown) and viable triploids were produced only at 41 and 41.4 °C. Viability and triploid yield was slightly in favor of HS at 41.4 °C, and thus it was used in the second HS trial assessing different HS durations. Fraction of detected triploids in HS treated embryos was significantly higher at HS lasting 1 and 2 min, and further HS prolongation resulted in significantly decreased survival rate only. Last HS trial tested different initiation time for HS treatment at 41.4 °C lasting 2 min. Treatment initiated 2 mpf (minutes post fertilization) yielded significant higher survival as well as triploid induction rate in comparison to 1mpf (Figure 1 D-F), moreover, embryos treated at 1mpf had less expanded chorion (Supplementary figure 2) and most of them did not hatched even when embryos appeared to be developed normally. Therefore, **treatment at 41.4 °C initiated 2 mpf and lasting 2 min was identified as optimal protocol for triploid induction and was used to produce recipients for germ cell transplantation in following experiment.**

**Figure 2.**
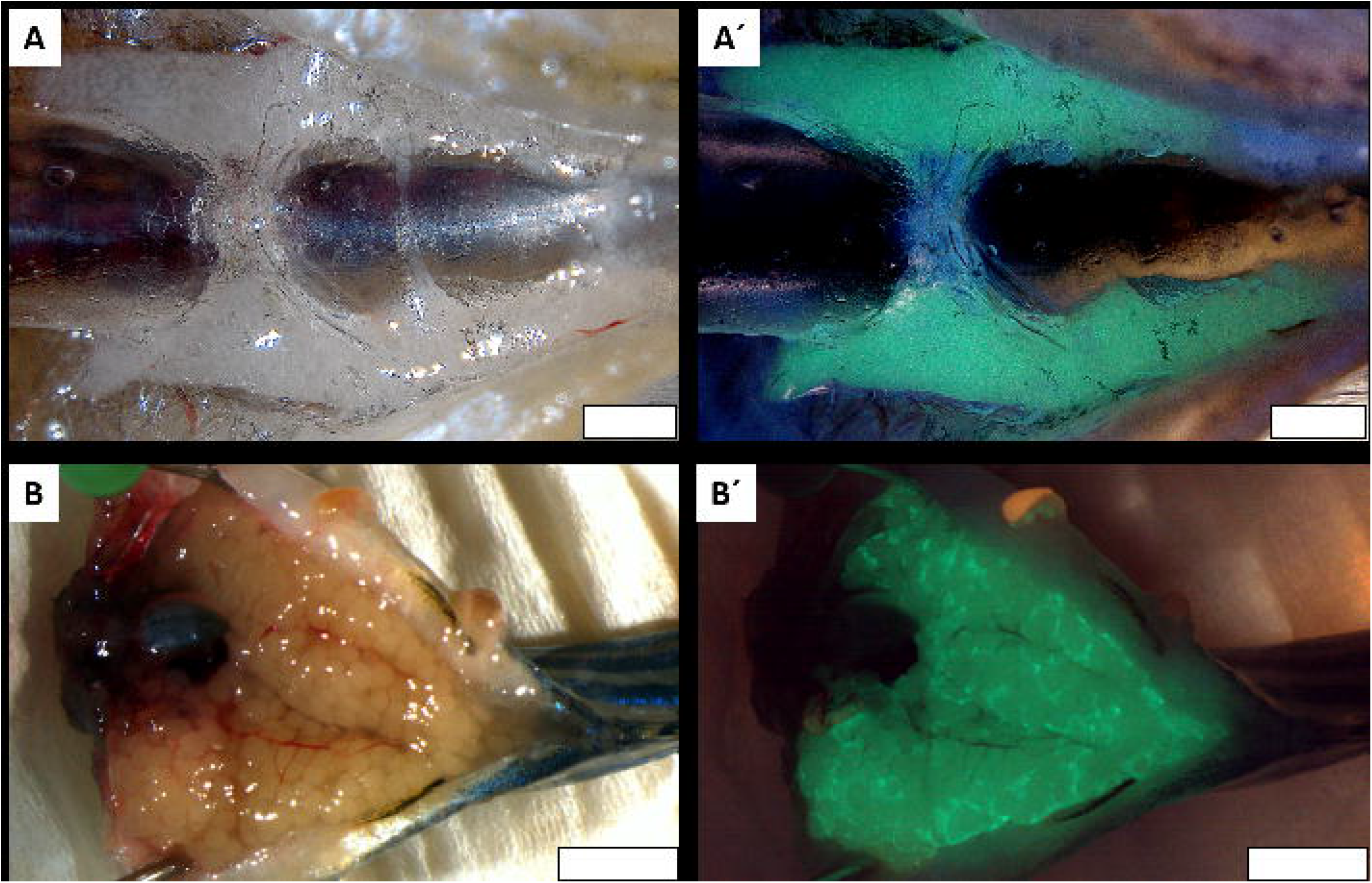
Donors used for germ cell transplantation into triploid surrogate recipients. Ventral view on dissected vas:EGFP donors, A) male, bright field and A’) fluorescent caption of testis, B) female, bright field and B’) fluorescent caption of ovaries. Scale bars A and A’ 2500 μm, B and B’ 5000 μm.

### 3.2 Germline chimera generation and reproduction

Despite of relatively invasive transplantation and sequentially presence of exogenous cells, the survival rate was similar among transplanted triploids, non-transplanted triploids and control diploids (Table 2). Transplanted testicular and ovarian cells from vas:EGFP donors (Figure 2) into triploid recipient showed strong signal after transplantation (Figure 3a). At 1wpt genital ridge of transplanted fish showed various patterns of germ cell colonization, large number of transplanted cells surrounded whole glass bladder (Figure 3 B); 5-20 cells located in the genital ridge close to the posterior part of the gas bladder; few individual cells located alongside the genital ridge. All patterns of colonization were represented in approximately same ratio. Transplanted ovarian cells were mostly found as a few or individual cells alongside the genital ridge, probably due to the lower number of transplanted cells. More than half of the positive germline chimeras receiving testicular cells showed the colonization bilaterally in the genital ridges. Noticeable proliferation of transplanted cells started at 7-12 dpt in majority of positive germline chimeras. Observation of transplanted fish at 2-3 wpt showed GFP positive cells proliferating and forming clusters alongside the gas bladder (Figure 3 C,E) or progression towards the anterior in case of cells originally colonizing posterior part of gas bladder. Observation of gonadal development was difficult due to deposits of fat cells surrounding gonads (Figure 3 E, E’). Typical patterns of vas:EGFP cells development in triploid recipients are shown in Figure 3. Fish were screened at 10 weeks post fertilization (wpf) for presence of GFP signal in gonads (Figure 4) and finclips of positive germline chimeras were taken for flow cytometry examination and triploidy of all positive chimeras was confirmed. All chimeras developed phenotypically in males regardless of origin of transplanted cells (testicular or ovarian). Colonization rate assessed by 10dpt was higher in TC groups in comparison to OC groups (Table 2).

**Figure 3.**
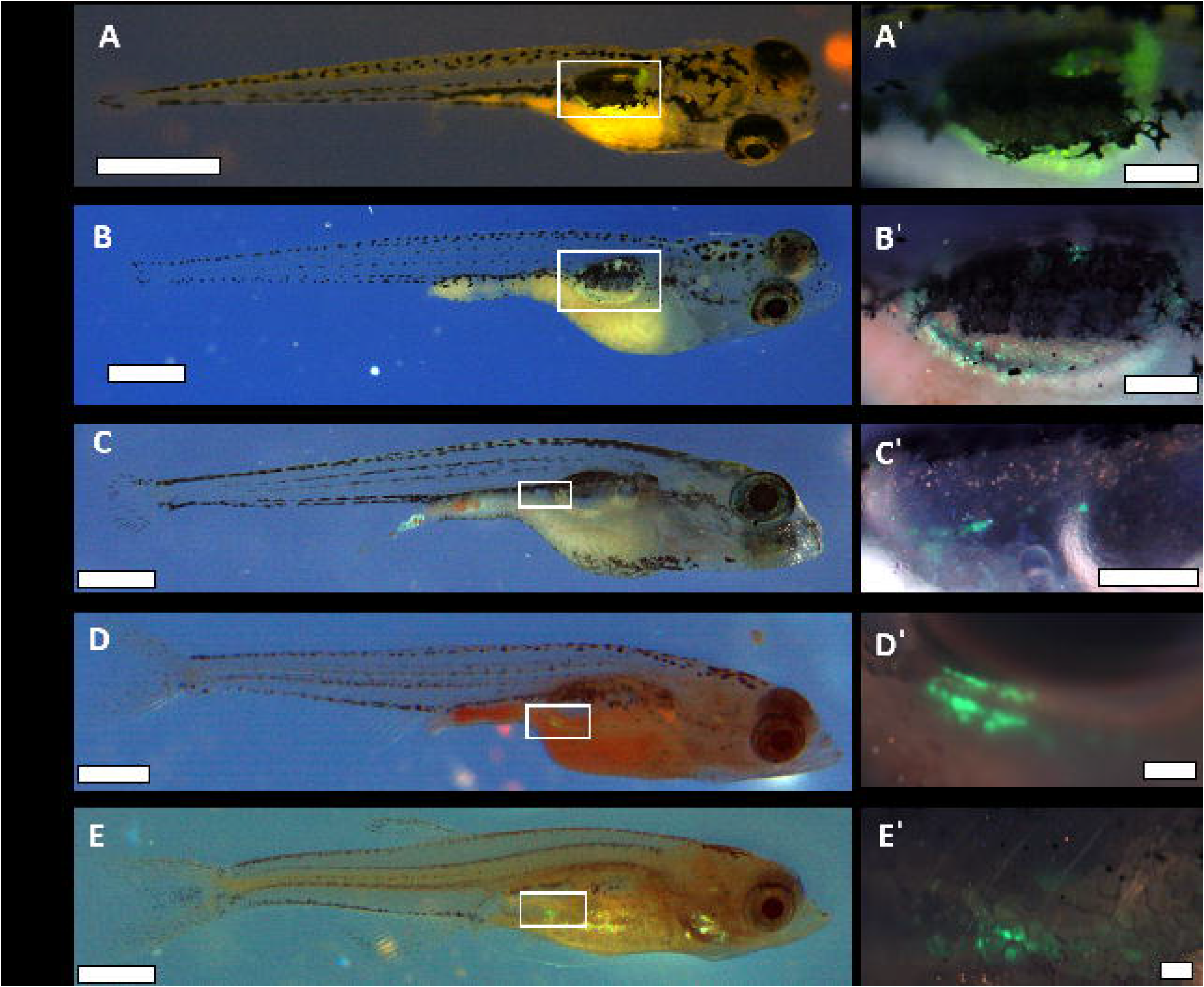
Patterns of colonization after vas:EGFP germ cells transplantation into triploid recipients. Colonization patterns were observed 24hpf until 4wpt. Captions were taken using DA/FI/TR fluorescent filter. A-E) view on whole fish, scale bar 2mm. White rectangle depicts magnified view on vas:EGFP transplanted cells A’-E’, scale bars 500μm.

**Table 2.** Overall results from testicular and ovarian germ cell transplantation into triploid zebrafish recipients

| Trial |  | I. |  |  |  | II. |  |  |  | III. |  |  |  |
| --- | --- | --- | --- | --- | --- | --- | --- | --- | --- | --- | --- | --- | --- |
| Group |  | TC | OC | 3n C | 2n C | TC | OC | 3n C | 2n C | TC | OC | 3n C | 2n C |
| Transplanted |  | 30 | 30 | 30 | 30 | 30 | 30 | 30 | 30 | 30 | 30 | 30 | 30 |
| 24hpt | Survival | 29 | 28 | 29 | 30 | 28 | 29 | 30 | 30 | 29 | 29 | 30 | 30 |
| 1wpt | Survival total | 27 | 26 | 28 | 29 | 28 | 26 | 28 | 28 | 27 | 26 | 26 | 27 |
|  | GFP + | 22 | 18 | - | - | 15 | 16 | - | - | 24 | 15 | - | - |
| 2wpt | Survival Total/GFP | 24/20 | 22/17 | 26/- | 28/- | 24/13 | 23/14 | 24/- | 26/- | 23/21 | 21/11 | 25/- | 26/- |
|  | GFP+ | 16 | 12 | - | - | 11 | 10 | - | - | 20 | 9 | - | - |
| 4wpt | Survival Total/GFP | 23/16 | 20/11 | 24/- | 27/- | 24/11 | 20/9 | 22/- | 26/- | 18/17 | 19/8 | 22/- | 26/- |
|  | GFP+ | 14 | 10 | - | - | 11 | 7 | - | - | 13 | 8 | - | - |
| 10wpt | Survival Total/GFP | 23/14* | 19/10* | 24/-* | 26/-* | 24/11 | 18/7 | 22/- | 26/- | 18/13 | 19/8 | 22/- | 23/- |
|  | GFP+ | 12 | 8 | - | - | 9 | 7 | - | - | 11 | 8 | - | - |
| Adult | Survival Total/GFP | 22/13 | 18/9 | 24/- | 26/- | 22/9 | 18/7 | 22/- | 26/- | 18/11 | 19/8 | 22/- | 23/- |
|  | GFP+ | 13 | 9 | - | - | 9 | 7 | - | - | 11 | 8 | - | - |
|  | GFP+ sperm | 11 | 7 | - | - | 7 | 5 | - | - | 9 | 7 | - | - |

Colonized part of the testis of dissected adults could be detected easily in bright filed according to its white color with observable lobules while control triploid testis were almost transparent (Figure 4). Exogenous cells were localized unilaterally mostly when middle and/or posterior part of the gonad was colonized with little extent towards anterior. Triploids developed phenotypic testis, with visible spermatogonia clusters, spermatids (including spermatids apparently in zygotene and pachytene stage) when only few spermatozoa were observed in the tubular lumens from nontransplanted control. Testis from control diploid testis had tubular lumens filled with spermatozoa. (Figure 5). Dissection showed positive GFP signal in triploid germline chimeras and in control donor vas:EGFP strain, while no GFP signal was detected in control non-transplanted triploid and normal control AB males.

**Figure 4.**
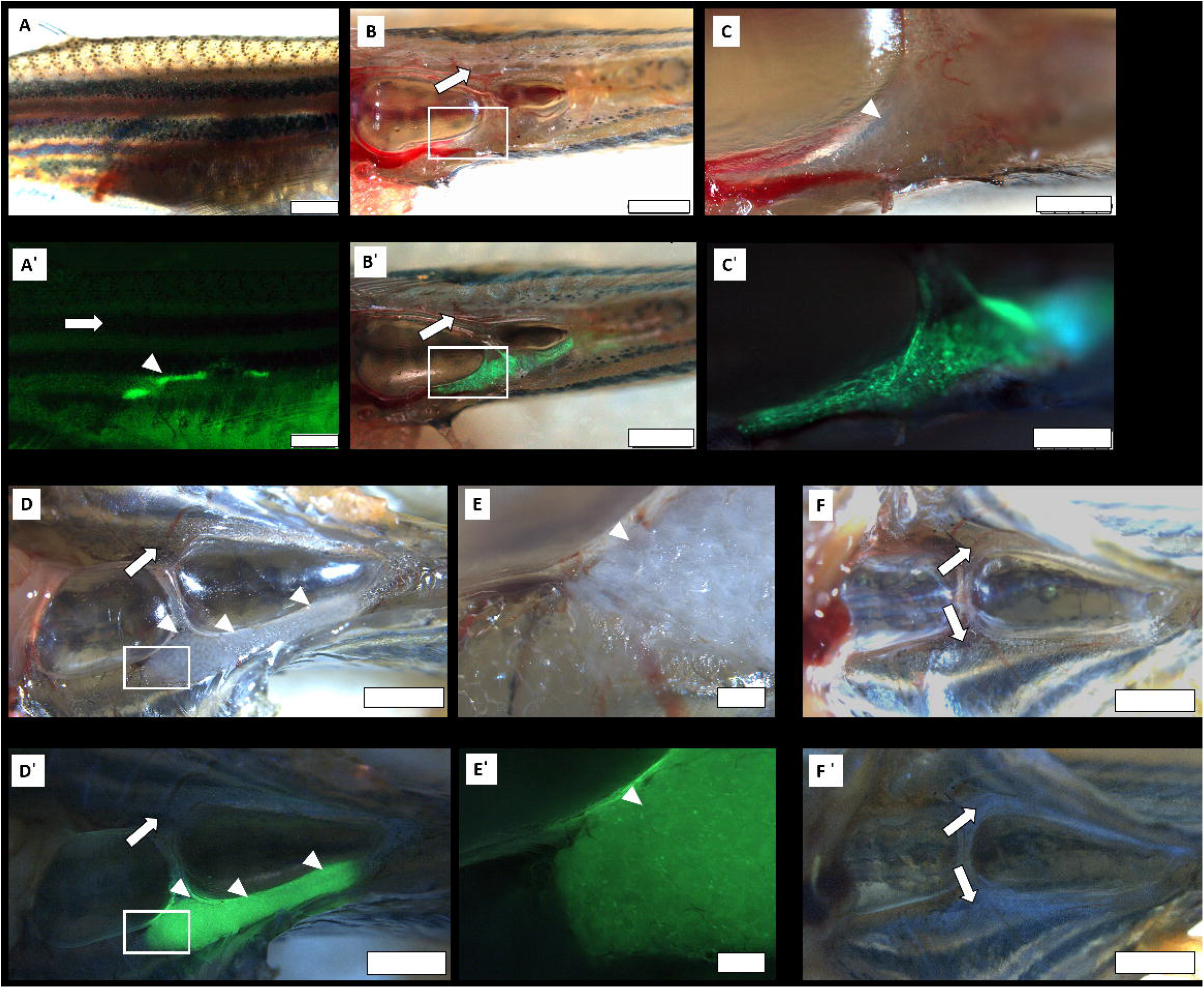
Gonadal development in juvenile and adult triploid germline chimera. A-B) observation of gonadal development at 10wpt, positive colonization could be detected *in vivo,* B’, C’) – germ cell colonization is detected by GFP signal and pointed out by white arrowhead. B’) ventral view on germline chimera gonads, with strong GFP signal in the right testis, left testis is non colonized (white arrow). White rectangles depict magnified captions of the colonized gonad (B’, C’ and E’). D, E’) ventral bright field and fluorescent view on gonads of adult triploid germline chimera. White rectangles depict magnified captions of the colonized gonad (C’-D’). White arrow indicates non colonized part of testis, white arrowheads indicate colonized testis with transplanted germ cells. F, F’) non transplanted triploid control. Scale bars A-B’, D-D’, F-F’ 2500 μm, C-C’ 750 μm, E-E’ 250 μm.

**Figure 5.**
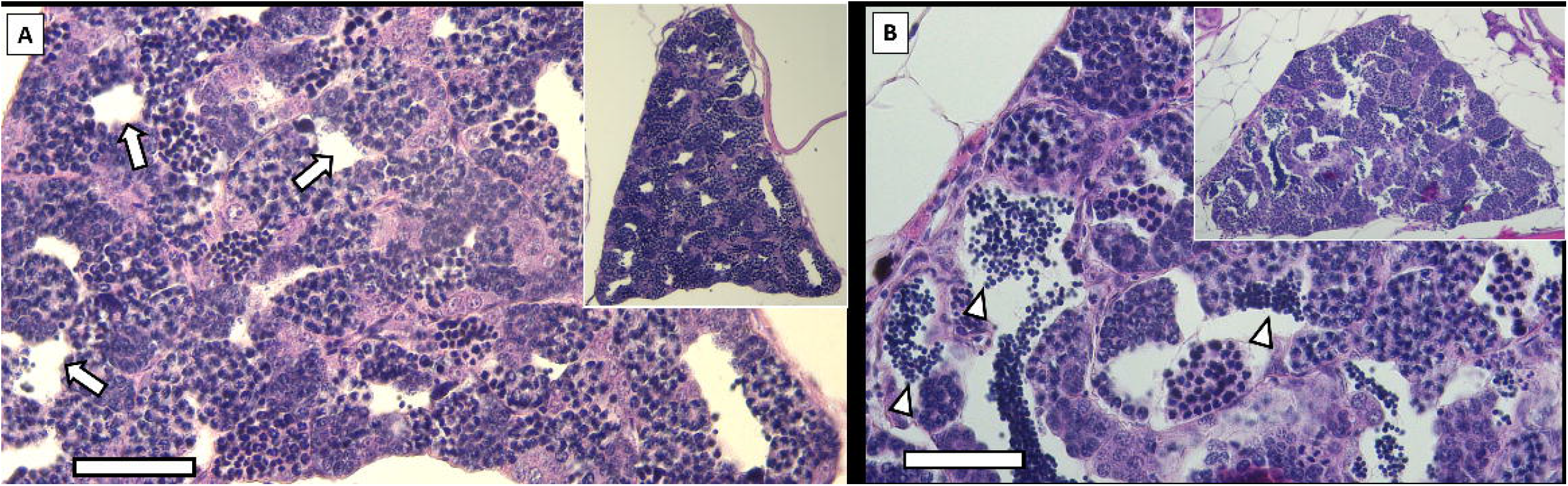
Photomicrographs of histological sections of zebrafish testis. View on whole testis is in right upper corner. A) triploid individual, empty lumen is pointed out by arrow. B) diploid control male, lumen with spermatozoon is pointed out by arrow. Scale bars 50 μm.

Table 2 shows over results of germ cell transplantation into triploid recipients. Success of the transplantation was evaluated as a total numbers of surviving fish until adulthood with detected positive GFP signal (GFP+) in their gonads evaluated *in vivo* and successful collection of GFP positive sperm from adult germline chimeras (GFP+ sperm). 3n C group represents remainder of the triploids from the batch used for transplantation. 2n C group is part of embryos non-treated by HS. From 1 wpt until adult whole group was always screened for positive GFP signal and subdivided into positive and negative group in order to be able to distinguish potential loss of signal from mortality. Survival Total/GFP represents number of fish surviving from previous screening counted before next screening. * Two GFP positive individuals from TC and OC and from 3n and 2n control groups were sacrificed for gonad observation.

Majority of GFP positive triploid germline chimeras produced sperm, with GFP signal detected in all collected samples (Table 2, Figure 6) and were capable to fertilize AB strain eggs during seminatural as well as *in vitro* fertilization. In overall, reproductive performance of triploid germline chimeras was similar to diploid control males from vas:EGFP strain, however, both tests showed that the control males from vas:EGFP always had the highest performance evaluated as fertilization rate, survival 24hpf and swim up rate, while germline chimeras transplanted by ovarian cell had the lowest survival rate (Table 2 and 3). Later PCR analysis confirmed 100% germline transmission, when GFP specific amplicon was detected (Table 3 and 4, Supplementary figure 4).

**Figure 6.**
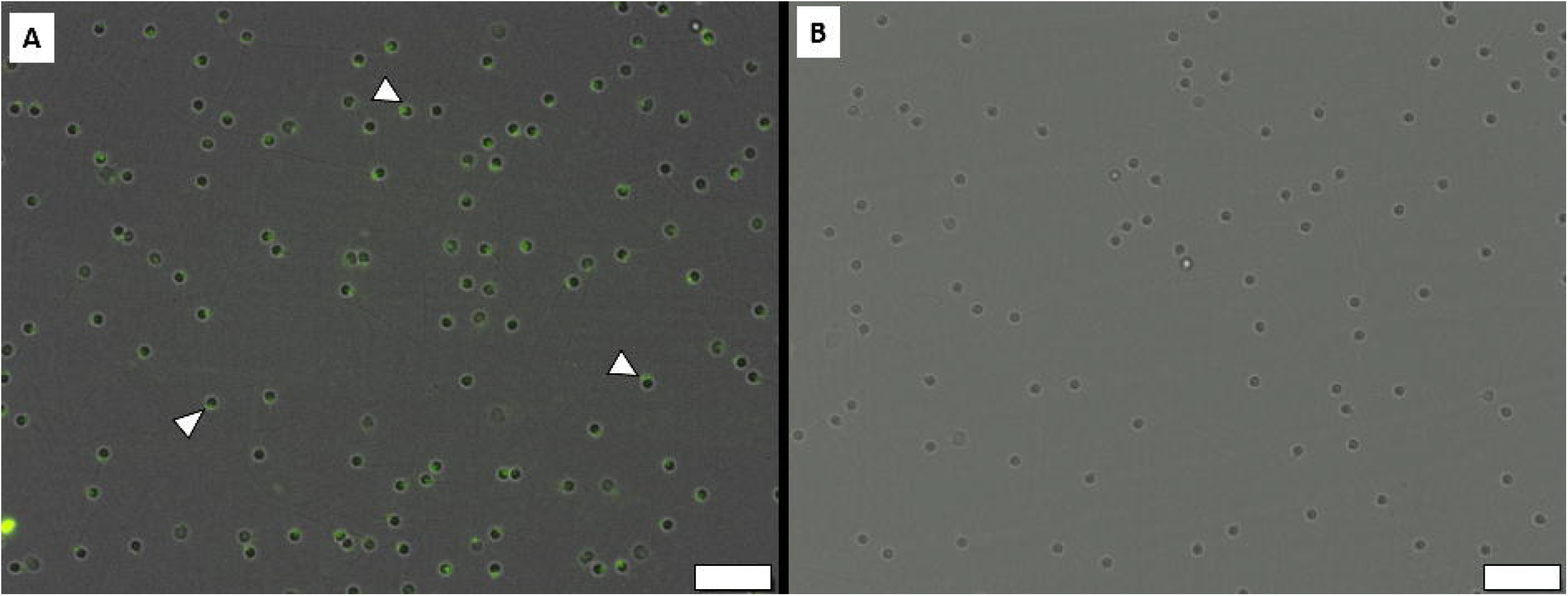
Analysis of germline transmission in triploid surrogates. A) fluorescent photomicrograph of sperm collected from triploid zebrafish germline chimera transplanted by testicular cells. White arrowheads indicate the head of spermatozoon with positive GFP signal. B) sperm collected from control diploid AB line male. Scale bars 20 μm.

**Table 3.** *In vitro* fertilization test of triploid germline chimeras producing donor-derived sperm

| Group | Male | Eggs | Fertilization rate | Survival 24hpf | Swim up rate | GFP - PCR |
| --- | --- | --- | --- | --- | --- | --- |
| Testicular cells | m1 | 39 | 32/82.1% | 25/64.1% | 21/53.8% | 10/10 |
|  | m2 | 66 | 60/90.9% | 55/83.3% | 49/74.2% |  |
|  | m3 | 85 | 65/76.5% | 58/68.2% | 51/60% |  |
|  | m4 | 52 | 46/88.5% | 40/76.9% | 34/65.4% |  |
|  | m5 | 64 | 54/84.5% | 47/73.4% | 37/57.8% |  |
|  | Σ | 306 | 84.5±5% | 73.2±6.7% | 62.3±7.1% |  |
| Ovarian cells | m1 | 56 | 46/82.1% | 46/82.1% | 42/75% | 10/10 |
|  | m2 | 41 | 35/85.4% | 31/75.6% | 18/43.9% |  |
|  | m3 | 87 | 84/96.6% | 72/82.8% | 59/67.8% |  |
|  | m4 | 67 | 49/73.1% | 41/61.2% | 32/47.8% |  |
|  | m5 | 43 | 31/72.1% | 25/58.1% | 15/34.9% |  |
|  | Σ | 294 | 81.9±8.9% | 72±10.4% | 53.9±15.1% |  |
| Control | m1 | 68 | 56/82.4% | 51/75% | 50/73.5% | 10/10 |
|  | m2 | 36 | 32/88.9% | 29/80.6% | 25/69.4% |  |
|  | m3 | 84 | 77/91.7% | 72/85.7% | 63/75% |  |
|  | m4 | 32 | 29/90.6% | 22/68.8% | 17/53.1% |  |
|  | m5 | 40 | 39/97.5% | 31/77.5% | 27/67.5% |  |
|  | Σ | 260 | 90.2±4.9% | 77.5±5.7% | 67.7±7.8% |  |

**Table 4.** Fertilization test of triploid germline chimeras after semi-artificial mating with AB females.

| Group | Male | Eggs | Fertilization rate | Survival 24hpf | Swim up rate | PCR - GFP |
| --- | --- | --- | --- | --- | --- | --- |
| Testicular cells | m1 | 149 | 82/55% | 73/49% | 69/46.3% | 10/10 |
|  | m2 | 76 | 55/72.4% | 48/63.2% | 32/42.1% |  |
|  | m3 | 134 | 82/61.2% | 70/52.2% | 62/46.3% |  |
|  | m4 | 52 | 38/73.1% | 35/67.3% | 28/53.8% |  |
|  | m5 | 82 | 51/62.2% | 44/53.7% | 34/41.5% |  |
| | $\Sigma$ | 493 | 64.8±6.9% | 57.1±7% | 46±4.4% | |
| Ovarian cells | m1 | 79 | 55/69.6% | 38/48.1% | 32/40.5% | 10/10 |
|  | m2 | 37 | 23/62.2% | 21/56.8% | 18/48.6% |  |
|  | m3 | 63 | 39/61.9% | 34/54% | 20/31.7% |  |
|  | m4 | 108 | 72/66.7% | 68/63.8% | 53/49.1% |  |
|  | m5 | 97 | 42/43.3% | 30/30.9% | 24/24.7% |  |
| | $\Sigma$ | 384 | 60.7±9.2% | 50.5±10.9% | 38.9±9.5% | |
| Control | m1 | 114 | 95/83.3% | 87/76.3% | 70/61.4% | 10/10 |
|  | m2 | 89 | 61/68.5% | 49/51.1% | 44/49.4% |  |
|  | m3 | 74 | 52/70.3% | 49/62.2% | 43/58.1% |  |
|  | m4 | 82 | 43/52.4% | 38/46.3% | 32/39% |  |
|  | m5 | 34 | 28/82.4% | 25/73.5% | 22/64.7% |  |
|  | m6 | 98 | 65/66.3% | 58/59.2% | 45/45.9% |  |
|  | m7 | 56 | 38/67.9% | 32/57.1 | 24/42.9% |  |
| | $\Sigma$ | 547 | 70.2±9.7% | 61.4±9.7% | 51.6±9.1% | |

Table 3 shows overall results of fertilization test when sperm collected from randomly chosen germline chimera males was used to fertilize pooled eggs obtained by stripping from four males. Fertilization rate, survival 24hpf and swim up rate is expressed in total numbers/%, summarized results from survival rates are expressed in % as mean ±S D. PCR – GFP shows results of detection of GFP specific amplicon in 10 randomly selected swim up larvae from pool in each group Table 4 shows overall results of fertilization test when germline chimeric males previously confirmed for GFP sperm production were randomly selected (10 males from each group) and set individually with two AB females and allowed to spawn. Note that only successful spawnings were included in this table, while five males from TC and OC, and three males from C group did not induce oviposition. Fertilization rate, survival 24hpf and swim up rate is expressed in total numbers/%, summarized results from survival rates are expressed in % as mean ± SD. PCR – GFP shows results of detection of GFP specific amplicon in 10 randomly selected swim up larvae from pool in each group

## 4 Discussion

Cold and heat shock treatments were tested in zebrafish in order to optimize method for triploid production. The produced triploids were then used as sterile recipients for surrogate reproduction. Heat shock treatment at 2 mpf, lasting 2min with temperature 41 °C was identified as the most suitable, for reliable triploid zebrafish production. All artificially induced triploids developed in phenotypic sterile males. We further tested their suitability as a surrogate parents when a fraction of testicular and ovarian cells was transplanted intraperitoneally. Colonization rates were in favour of testicular cells, however, only male triploid germline chimeras, which were fertile and capable to mate with females from AB strain, were produced.

### 4.1 Triploid induction

First triploid induction in zebrafish have been reported by [33], when fertilized eggs were treated at 2.5 mpf at 41 °C, for 4 min, however, this conditions always resulted in complete mortality in our conditions. Other studies used abovementioned protocol with slight modification such as at 2.5 mpf at 41 °C, for 2 min [36] or 2 mpf at 41 °C, for 2 min [37]. Our results showed, that only heat shock treatment is suitable for effective triploid production. Only partial fraction of triploid swim up larvae was obtained after optimized cold shock treatment, moreover, survival was significantly higher after HS compared to CS. All adult triploids developed in phenotypic males with testis almost free of spermatozoon, while control diploid males showed lumens filled with spermatozoon. Apparently large proportion of germ cells in triploid testis were observed to be arrested in pachytene of the first meiosis as it is result of odd chromosome number exhibiting in disorganized synapsis [38]. These results confirmed previously reported findings when all triploid zebrafish with some exception where few female individuals among males. In our study, no triploid females were detected at all. Only male occurrence in artificially induced triploid is extremely rare in fish, to the best of our knowledge documented in zebrafish [37] and Rosy bitterling [39].

### 4.2 Surrogate reproduction

Artificially induced triploids have been used successfully as recipients for surrogate gamete production in several fish species such as masu salmon [2], grass puffer [14], medaka [9], rainbow trout [40,41] and nibe croaker [13]. To the best of our knowledge this study provided the first report of suitability suitability of triploid swim up larvae as a recipient for intraspecific transfer of germ cells and production of donor-derived gametes. As was previously described, triploid zebrafish developed in males only [37], even after rescuing their fertility by transplantation of testicular or ovarian cells.

Germ stem cells have been proved to be bipotential gamete precursors as they can develop in recipients gonads into female or male germ cells according to the recipient’s sex [3]. Spermatogonia transplantation in species with male heterogamety resulted in partial production of YY rainbow trout supermales after mating male and female germline chimeras. This approaches could serve as an alternative for mono sex culture production which is normally achieved by production and subsequent mating of androgenetic or gynogenetic stocks [42].

Situation in zebrafish and sex control is more complicated, since some families can produce extremely biased offspring when percentage of males can vary from 4.8% to 97.3% [43] or from 0% to 75% when fish were challenged to unfavorable or affluent conditions [44]. This phenomenon is attributed to polygenic sex determination with the further influence of the surrounding environment [45,46], moreover, two zebrafish lines have been shown to lack sex-linked loci [47]. Thus, a different subpopulation of zebrafish can produce progeny in very fluctuating sex ratios.

Theoretically, part of progeny produced using sperm from triploid germline chimeras transplanted by ovarian cells should after fertilization of normal eggs should yield fraction of WW super females progeny, which could be interesting model for other fish species possessing female heterogamety sex determination. Then in turn, induction of triploidy with of eggs obtained from WW females fertilized with sperm from triploid germline chimera possessing W or Z chromosome should yield fraction of WWW super female triploids, which could provide more insights into sex determination in zebrafish and only triploid male occurrence.

Real use of zebrafish recipients in surrogate reproduction resulted in only male germ line chimera production independently on used approach of zebrafish sterilization and type of germline transfer such as using blastomeres, single PGCs or adult germ stem cell [15,48,49]. Production fertile zebrafish female chimera seems to be not possible so far. Reason for literally absence of germline chimera females is attributed to sterilization of recipient by PGCs depletion. In zebrafish, certain numbers of PGCs are required to maintain the ovarian fate [17]. When taking in account that very few of transplanted cells are capable to colonize the recipient’s gonad, it is clear that such low number of cells bellow a threshold (3-29 PGCs) cannot maintain ovarian fate. Moreover, it has been shown that female germ cell presence is essential even in adulthood to maintain ovarian fate and prevent sex reverse into functional male [50].

In conclusion to carry out whether and how to produce zebrafish germline chimeras producing eggs, following possibilities have not been tested yet. 1) Hormonal treatment optimization for zebrafish germline chimeras as was firstly attempted by Saito et al., (2008) on zebrafish x pearl danio hybrid when 3 from 4 fish developed as females, but were not able to produce eggs. 2)

Increasing a number of germ cells colonizing the recipient gonad might have an effect on sex differentiation in germ line chimera as was proven for a number of PGCs, since so far it was shown that only few individual cells are colonizing gonads after transplantation. 3) Cotransplantation of female germ stem cells with early oocytes could also act supportively for female sex differentiation in germline chimera, however, this method has not been tested yet. 4) Essentiality of *dmrt1* and *amh* gene for proper male development have been reported recently in zebrafish [51,52], thus DNA and RNA interfering approaches such knockdown or knock out could influence sex ratio in germline chimeras in favour of females.

## 5 Conclusion

Surrogate reproduction via germ cell transplantation into zebrafish triploid developed in this study could have potential to serve as an alternative way for zebrafish gene resource banking since it can be combined with a convenient method for spermatogonia cryopreservation by needle immersed vitrification [53]. Up to day, literally, thousands of mutants, transgenic lines and recently CRISPR/Cas9, ZFN or TALEN genetically engineered strains have been generated in zebrafish which makes gene banking of utmost importance [54–56]. Similarly, triploid males can be used as recipients to improve sperm production when originally few individuals are available for breeding or given line suffer from poor reproductive performance as was shown in medaka when the reproductive performance of an inbred strain was improved by transplantation into triploid surrogates [9]. From our experience, a number of early-stage germ cells obtained from testis originating from single adult zebrafish male is sufficient for intraperitoneal transplantation into at least 40-50 individuals. Thus taking into account that at least 23 % of transplanted triploid zebrafish produced donor sperm (TC group), at least 10 fertile triploid males can be recovered using testis from a single donor. It is noteworthy to point out that triploid zebrafish germ line chimeras in our study were capable to mate with females from AB line in common spawning chamber, and their reproductive characteristics were comparable to mating with normal diploid males. Described HS protocol for triploid production is a simple method for sterile zebrafish production which does not require microinjection in embryos for delivery of compounds for gene knockdown or knock out to ensure sterilization. However, only sperm production from using PGCs depleted or hybrid recipient leaves an issue which needs to be addressed in order to produce donor derive eggs from zebrafish recipient.

## Acknowledgements

Republic, projects CENAKVA (LM2018099) and Biodiversity (CZ.02.1.01/0.0/0.0/16_025/0007370), by the Grant Agency of the University of South Bohemia in České Budějovice GAJU 097/2019/Z and 034/2017/Z. The Czech Science Foundation (project No. 17-19714Y) and NAZV QK1910248.

## Contribution and disclosure

RF and MP: conceptualization, designing of the study, performing experiments, data collection and funding acquisition, TT: data collection and analysis, CS and MF: ploidy analysis and histology sections. All authors contributed on manuscript drafting and approved the submitted version.

## Declarations of interest: none

The funders had no role in study design, data collection and analysis, and preparation of the manuscript.

