## Supplementary material for "Production and use of triploid zebrafish for surrogate reproduction": SI captions and figures

**Supplementary files**

Supplementary figure 1

Occurrence of deformities after heat shock treatment

A) Normally developing tripoid individual, B) heart edema, C) heart and body cavity edema, D) malformed embryo without developed eyes, E) malformed embryo incapable to hatch.

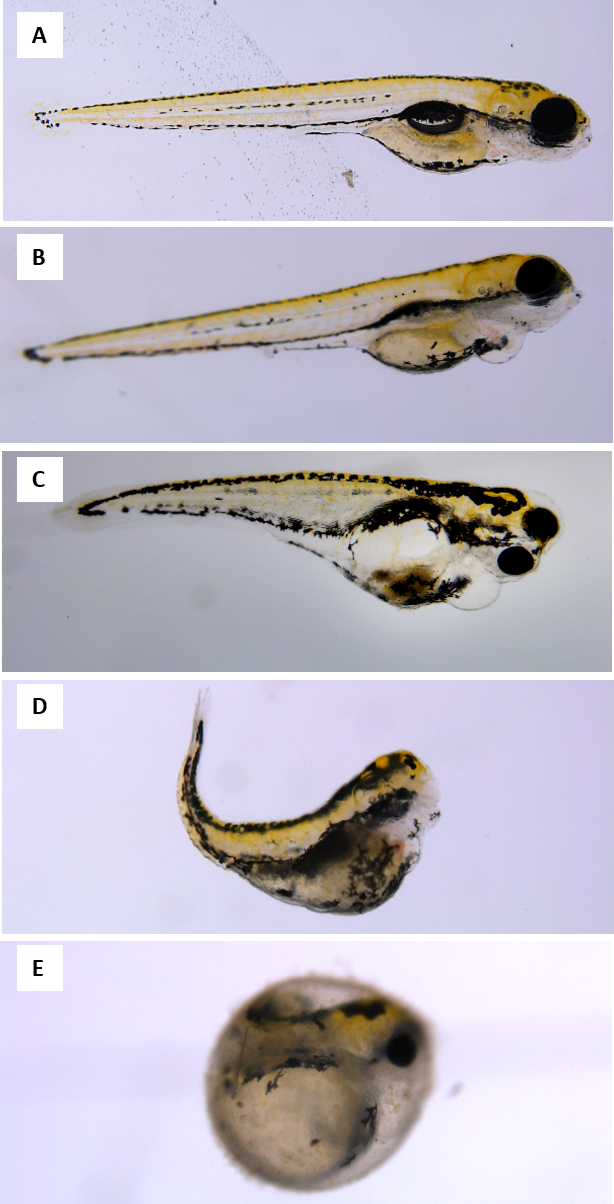

Supplementary figure 2

Swollen and non-swollen chorion in zebrafish embryos after heat shock treatment. Upper left embryo – heat shock starting at 2 min post fertilization, upper right embryo – untreated control. Lower embryos – heat shock starting at 1 min post fertilization. Scale bar 1 mm.

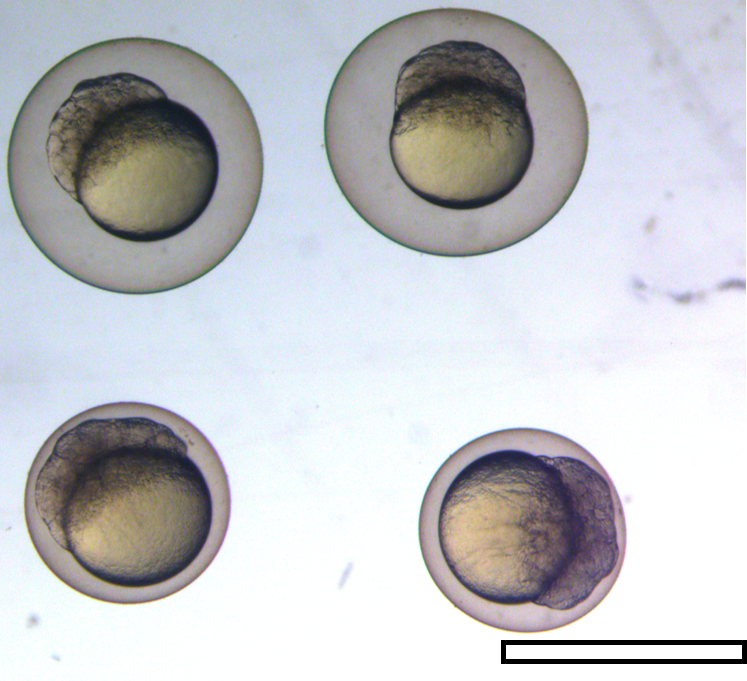

Supplementary figure 3

Results of flow cytometry analysis of swim up embryos after triploidy induction. A) Individual from heat shock group showing 1.5x higher relative DNA content corresponding to triploidy in comparison to B) control diploid.

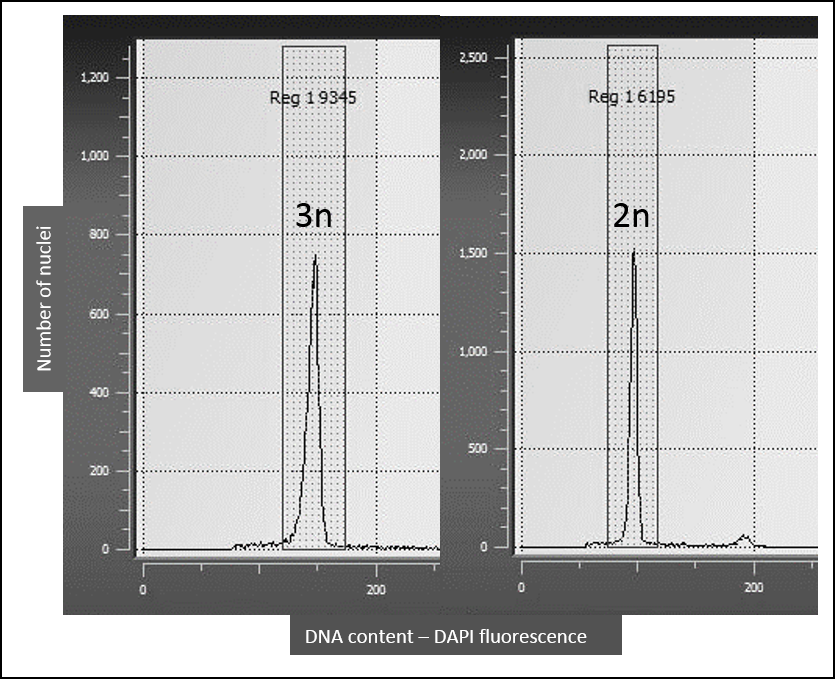

Supplementary figure 4

Detection of GFP specific amplicons in eight individuals from mating between germline chimera and AB females after PCR with GFP specific primers.

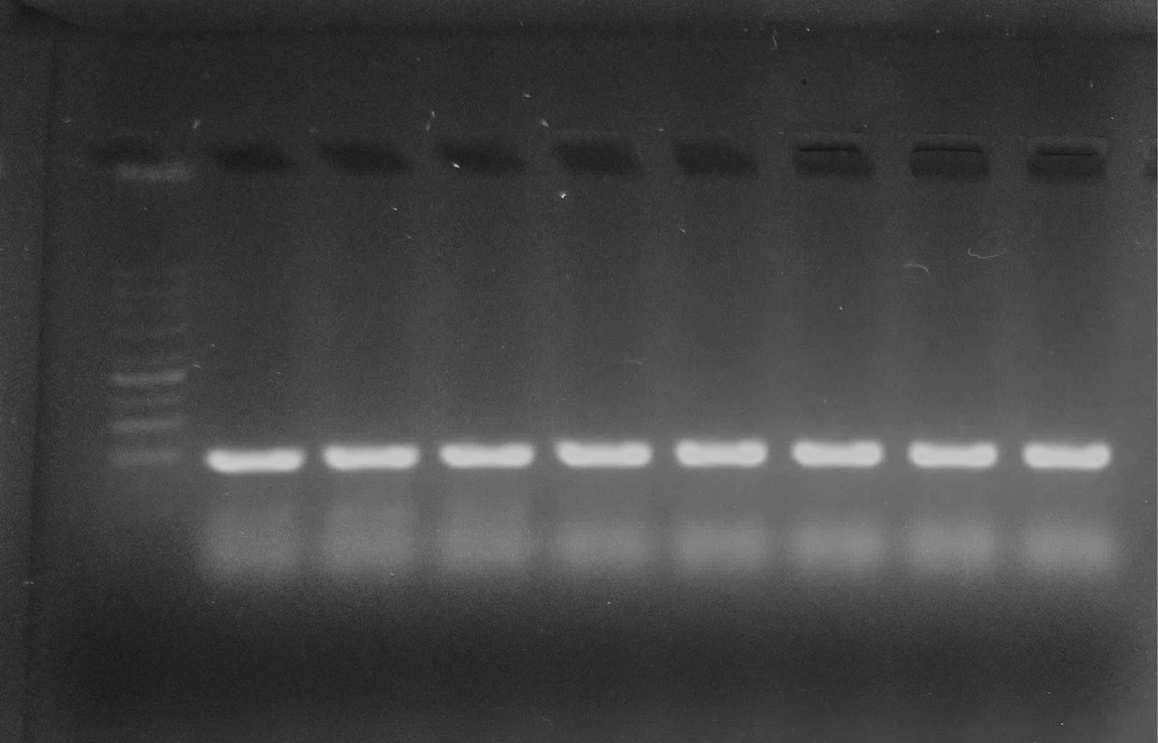
